# Gluconeogenesis in Aqueous Microdroplets: Non-Enzymatic Generation of Glucose

**DOI:** 10.1101/2025.06.17.660170

**Authors:** Jaeho Ko, Yukyung Kim, Jeongcheol Lee, Jae Kyoo Lee

## Abstract

Glucose is a central metabolite of living organisms, serving as the primary energy substrate produced predominantly by photosynthetic organisms. Under conditions of limited external glucose supply, organisms activate gluconeogenesis, an endogenous biosynthetic pathway that sustains essential glucose levels. Yet the mechanism of glucose generation, before the advent of photosynthesis and complex enzymatic systems, remains elusive. Recently, microdroplet chemistry has emerged as a novel approach for catalyst-free organic synthesis. Here, we report the non-enzymatic formation of glucose from a simpler organic precursor, pyruvate, in aqueous microdroplets without the aid of organic or inorganic catalysts. Our results show that glucose is generated via a reaction pathway analogous to canonical gluconeogenesis, proceeding through key intermediates including oxaloacetate, glycerate, and glyceraldehyde. Furthermore, thermodynamic analysis indicates that the free energy change associated with glucose formation is overcome in aqueous microdroplets at room temperature, without the need for external energy input or enzymatic catalysis. These findings indicate that aqueous microdroplets can non-enzymatically convert C_3_ compound, pyruvate, into the C_6_ sugar glucose, offering a plausible abiotic route for anabolic carbon transformations during abiogenesis.

## Introduction

Glucose (GC) is a critical monosaccharide that serves as a primary energy carrier and a key metabolic carbon source. Cellular activities including cellular respiration, glycolysis, and oxidative phosphorylation rely on glucose^1,2^. In addition, glucose is essential for synthesizing cellulose, the primary component of plant cell walls as well as other cellular membrane constituents^3^. In nature, glucose is predominantly produced through photosynthesis in plants and algae, whereas heterotrophic organisms obtain the glucose by consuming these external sources. When external glucose source is limited or intracellular levels decline, organisms activate endogenous glucose synthesis pathways, collectively known as gluconeogenesis, to maintain functional integrity^4,5^. Gluconeogenesis is a complex anabolic pathway involving a series of enzyme-catalyzed reactions and multiple intermediates with ATP serving as an energy source. This unique pathway generates glucose from non-carbohydrate precursors, such as pyruvate^4^. Moreover, gluconeogenesis is highly conserved across life, even in primitive organisms such as bacteria^6-8^ and archaea^7^.

Recent studies have demonstrated that key steps of central metabolic cycles, including the rTCA cycle^9^, tricarboxylic acid (TCA) cycle^10,11^, and Calvin cycle^12^, can be reconstructed chemically in the absence of enzymes. These findings suggest that fundamental metabolic pathways may function under abiotic conditions. Gluconeogenesis, a biosynthetic pathway that produces glucose from pyruvate, is significantly more complex. Its dependence on multiple enzymes operating in a highly coordinated sequence has so far prevented its realization in any abiotic setting. Instead, abiotic sugar synthesis has primarily been investigated through chemical routes^13-25^. Among these, the formose reaction is the most extensively studied. In this process, formaldehyde undergoes successive aldol condensation and isomerization steps to form various sugar compounds^13^. This reaction, however, produces a mixture of sugars—including pentoses and hexoses ^26,27^, and typically requires specific catalysts, such as CaCO_3_^22^, Mg(OH)_2_^23^, BaCl_2_, KOH^24,25^, and thiazolium^25^, under high-temperature and high-pressure conditions^13^. Thus far, numerous attempts have been made to synthesize monosaccharides. Despite numerous efforts, the abiotic generation of glucose from simple biological compounds without any enzymatic or catalytic machinery under ambient conditions remains challenging.

Recently, the non-catalytic synthesis of compounds using microdroplets has gained attention due to the distinct physical and chemical properties of these systems. Reactions in microdroplets proceed at accelerated rates compared to bulk-phase reactions owing to localization of molecules at the microdroplet interface^28^, the presence of strong interfacial electric fields ^29^, and extreme pH environments at the air-water interface^30^. These conditions create unique microenvironments that concentrate reactants, stabilize transition states, and enable otherwise inaccessible chemical transformations^31^. The confined microenvironments within microdroplets can concentrate reactants, stabilize transition states, and enable reaction pathways that are unavailable in bulk solutions^29^. These properties have enabled fundamental chemical transformations relevant to biomolecule synthesis. Reactions such as carboxylation^32^, reduction^33^, oxidation^34^, hydrogenation^35^, and phosphorylation^36^ have been successfully demonstrated in microdroplets. In addition, microdroplet synthesis systems have been shown to synthesize biologically important compounds, including peptides^37,38^, phospholipids^39^, and other organic polymers without the need for external catalysts^40^. These compounds play central roles in metabolic networks. Thus, the distinctive physicochemical characteristics of microdroplets make them valuable systems for the abiotic synthesis of biologically relevant molecules.

Here, we investigate the abiotic synthesis of glucose by reconstructing the gluconeogenesis pathway from the simple C3 compound pyruvate using microdroplet chemistry. Pyruvate was selected as the reactant due to its central role as a precursor in gluconeogenesis. It is also one of the most abundant and chemically simple carbon molecules, making it an ideal candidate for glucose synthesis experiments. We utilize aqueous microdroplets to achieve the non-enzymatic, non-catalytic conversion of pyruvate to glucose at room temperature, without the addition of ATP or other external energy sources. **Figure 1** presents the overall reaction scheme for glucose formation from pyruvate in aqueous microdroplets. Green arrows indicate the canonical biological gluconeogenesis pathway, while red arrows denote the reaction steps observed in aqueous microdroplets. Whereas intermediates in biological gluconeogenesis are typically phosphorylated, our system uses unmodified carbon-based compounds as reactants, since microdroplets can drive thermodynamically unfavorable reactions without enzymatic catalysis or ATP input^31^. The generation of glucose and intermediates, including oxaloacetate (OAA), glycerate (GA), and glyceraldehyde (GAH), in aqueous microdroplets reactions is confirmed. By systematically varying the microdroplet conditions, we identify factors that enhance non-enzymatic glucose production. The thermodynamic behavior of these reactions is further examined to elucidate plausible reaction mechanisms. These findings advance our understanding of abiotic sugar synthesis pathways and highlight aqueous microdroplets as chemically active microenvironments capable of facilitating non-enzymatic, multi-step reaction pathways traditionally considered inaccessible without biological machinery.

**Fig. 1.**
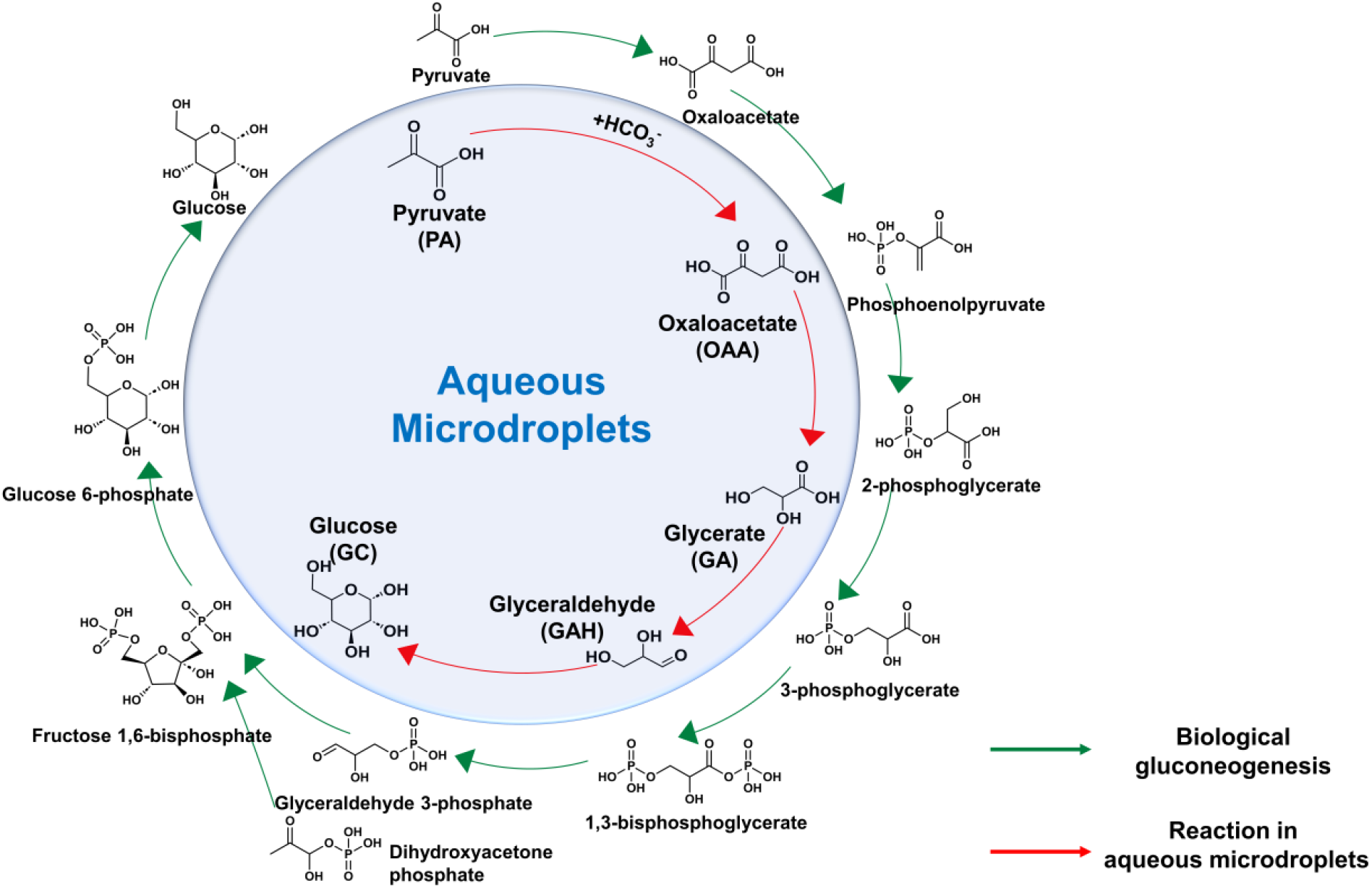
Schematic illustration of the reaction pathway for glucose formation from pyruvate in aqueous microdroplets (red arrows) and the canonical biological gluconeogenesis pathway (green arrows).

## Results

### Generation of glucose and gluconeogenetic intermediates in aqueous microdroplets

**Fig. 2a** illustrates the experimental setup for microdroplet reactions and mass spectrometric analysis of the products. Aqueous microdroplets were sprayed into a mass spectrometer (MS) inlet by nebulizing a solution containing reactants with dry N_2_ gas at a pressure of 0.8 MPa. A voltage of -5 kV was applied. Droplet diameters ranged from 1 to 50 μm^41^. The travelling distance from the capillary tip to the MS inlet was set to approximately 15 mm. The corresponding travel time of microdroplets at that distance was approximately 214 μs ^42,43^. All experiments were conducted at atmospheric pressure and room temperature. This experimental configuration is widely used in microdroplet chemistry studies^29,44-46^. **Fig. 2b** shows the setup for generating microdroplets using a vibrating mesh, which was used for ^1^H NMR analysis **(Fig. 2d)** and the glucose assay **(Fig. 3e)**. The reaction distance was set to approximately 10 cm. The average droplet diameter was 5 ± 0.5 µm, with a flow rate of 0.8 mL/min. No external voltage was applied during the droplet generation to avoid the potential influence of high voltage on glucose formation.

**Fig. 2.**
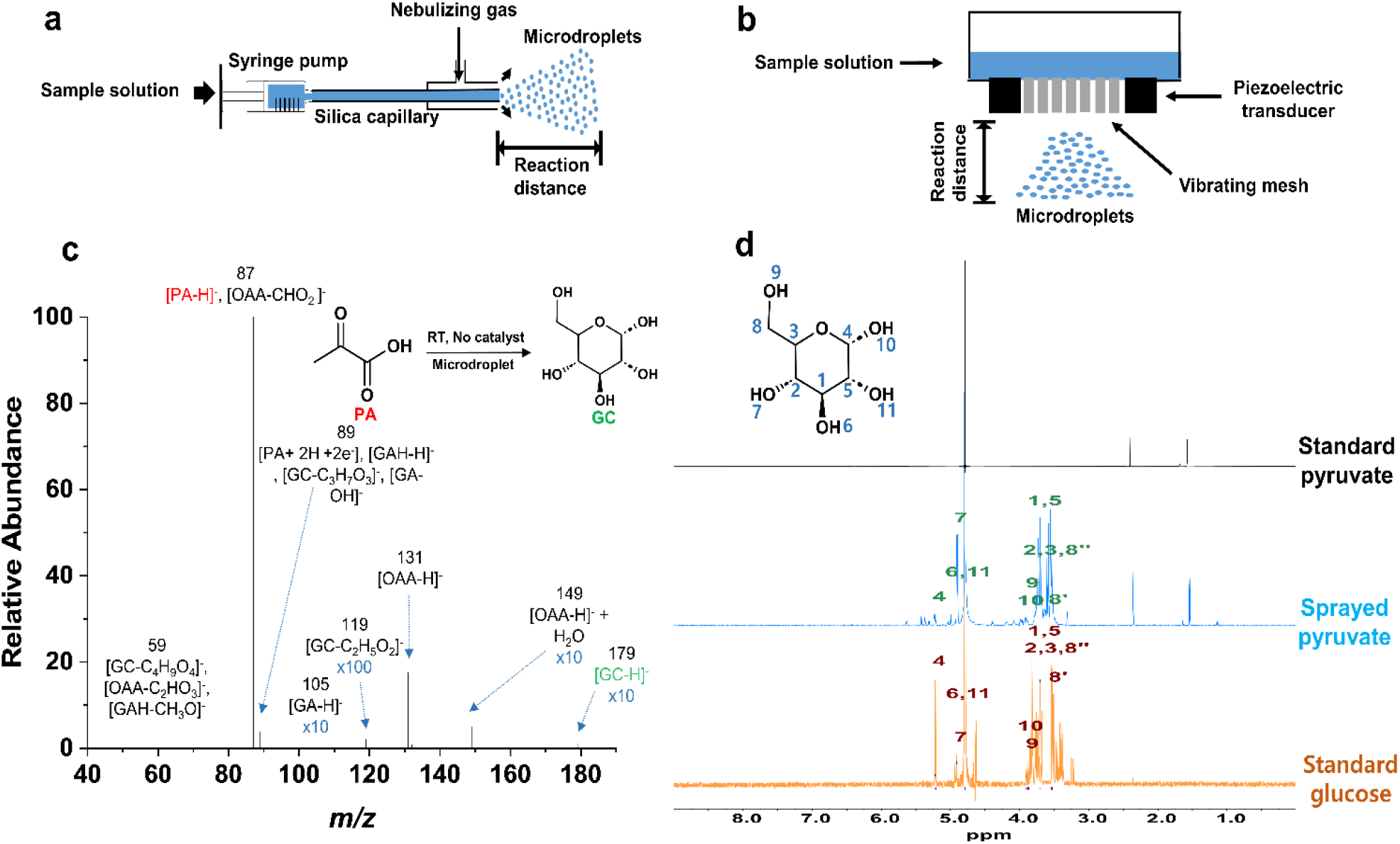
Generation of glucose and gluconeogenic intermediates from pyruvate in aqueous microdroplets. **a** Schematic of the experimental setup for mass spectrometry analysis. A silica capillary with an inner diameter of 50 µm and an outer diameter of 360 µm was used. A voltage of ±5 kV was applied. **b** Microdroplet generation setup using a vibrating mesh aerosolizer. Droplets with a diameter of 5 ± 0.5 µm were generated through mesh apertures. **c** Negative ion mode mass spectrum of aqueous microdroplets containing 10 µM pyruvate, acquired at a reaction distance of 15 mm. Intermediates, including oxaloacetate (OAA), glycerate (GA), and glyceraldehyde (GAH), as well as the product, glucose (GC), were detected. **d** ^1^H NMR spectra of standard pyruvate (black), sprayed pyruvate (blue), and standard glucose (orange), each at 10 mM in D_2_O.

We first investigated the formation of glucose from pyruvate in aqueous microdroplets. An aqueous solution containing 10 μM pyruvate was nebulized into the MS inlet. The negative ion mode mass spectrum of the sprayed solution exhibited a peak corresponding to glucose (*m/z* 179.05642, 1.73 ppm) **(Fig. 2c)**. To confirm the identity of this product, we performed higher-energy collisional dissociation (HCD) tandem mass spectrometry analysis. **Supplementary Fig. 1a, b** present the tandem MS/MS spectra of standard glucose and the compound detected at *m/z* 179 from the sprayed microdroplets, confirming the presence of glucose. The mass spectrum also displayed other peaks corresponding to known gluconeogenic intermediates: oxaloacetate (*m/z* 130.99898, 2.90 ppm), glycerate (*m/z* 105.01964, 2.95 ppm), and glyceraldehyde (*m/z* 89.02462, 2.25 ppm) **(Fig. 2c)**. Tandem MS/MS spectra of *m/z* 131, 105, 89 revealed fragmentation patterns consistent with those of standard oxaloacetate **(Supplementary Fig. 2)**, glycerate **(Supplementary Fig. 3)**, and glyceraldehyde **(Supplementary Fig. 4)**, confirming their molecular identities.

In biological gluconeogenesis, pathway intermediates typically contain phosphate groups that function as electron donors^47^. In contrast, our experiments were conducted in the absence of phosphate, yet yielded intermediates structurally analogous to those in the canonical gluconeogenesis pathway. These observations indicate that the reactions occurring in aqueous microdroplets proceed through pathways that resemble biological gluconeogenesis despite the lack of enzymatic or phosphorylated intermediates. Additionally, conventional gluconeogenesis involves fructose as an intermediate. Fructose and glucose are structural isomers that share the same chemical formula C_6_H_12_O_6_. To determine whether fructose was also generated, we compared the MS/MS spectrum of standard fructose with that of the molecular species observed at *m/z* 179 in sprayed solution, revealing no evidence of fructose formation in the microdroplet system. **(Supplementary Fig. 5)**.

The identity of glucose produced in aqueous microdroplets was further verified with ^1^H NMR spectroscopy. **Fig. 2d** shows ^1^H NMR spectra of standard pyruvate (black), sprayed pyruvate (blue), and standard glucose (orange), with all samples prepared at 10 mM in D_2_O. Key proton resonances are annotated, and complete spectral assignments, including chemical shifts (δ), multiplicity, and coupling constants (J), are provided in the Supplementary Information. Notably, ^1^H NMR signals corresponding to glucose were observed only in microdroplets generated from the nebulized pyruvate solution and were not observed in the bulk pyruvate solution. Taken together, these results demonstrate that pyruvate can undergo abiotic conversion to glucose in aqueous microdroplets.

Subsequently, intermediates of microdroplet reactions were further investigated. To confirm the involvement of oxaloacetate, glycerate, and glyceraldehyde as reaction intermediates in aqueous microdroplets, the generation of glucose from these compounds in microdroplets was evaluated. Aqueous solutions containing each intermediate prepared at 10 μM were nebulized into the MS inlet **(Fig. 3a)**. The resulting mass spectra showed the formation of glucose **(Fig. 3b-d)**. The molecular species at *m/z* 179 from the sprayed microdroplets of these intermediates exhibited MS/MS fragmentation patterns consistent with those of standard glucose **(Supplementary Fig. 1a, 1c-e)**.

To further validate glucose formation from different intermediates and to quantify glucose yields in aqueous microdroplets, the Picoprobe Glucose Assay Kit was employed **(Fig. 3e)**. The glucose assay, based on glucose oxidase, is a widely used method for quantifying glucose concentrations ^48^. All microdroplet samples (10 μM) were collected using the aerosol generator **(Fig. 2b)**. Glucose oxidase exhibits high specificity for D-glucose due to the stereoselectivity of its active site ^49^. Accordingly, the glucose detected in our experiments is primarily in the D-glucose form.

**Fig. 3.**
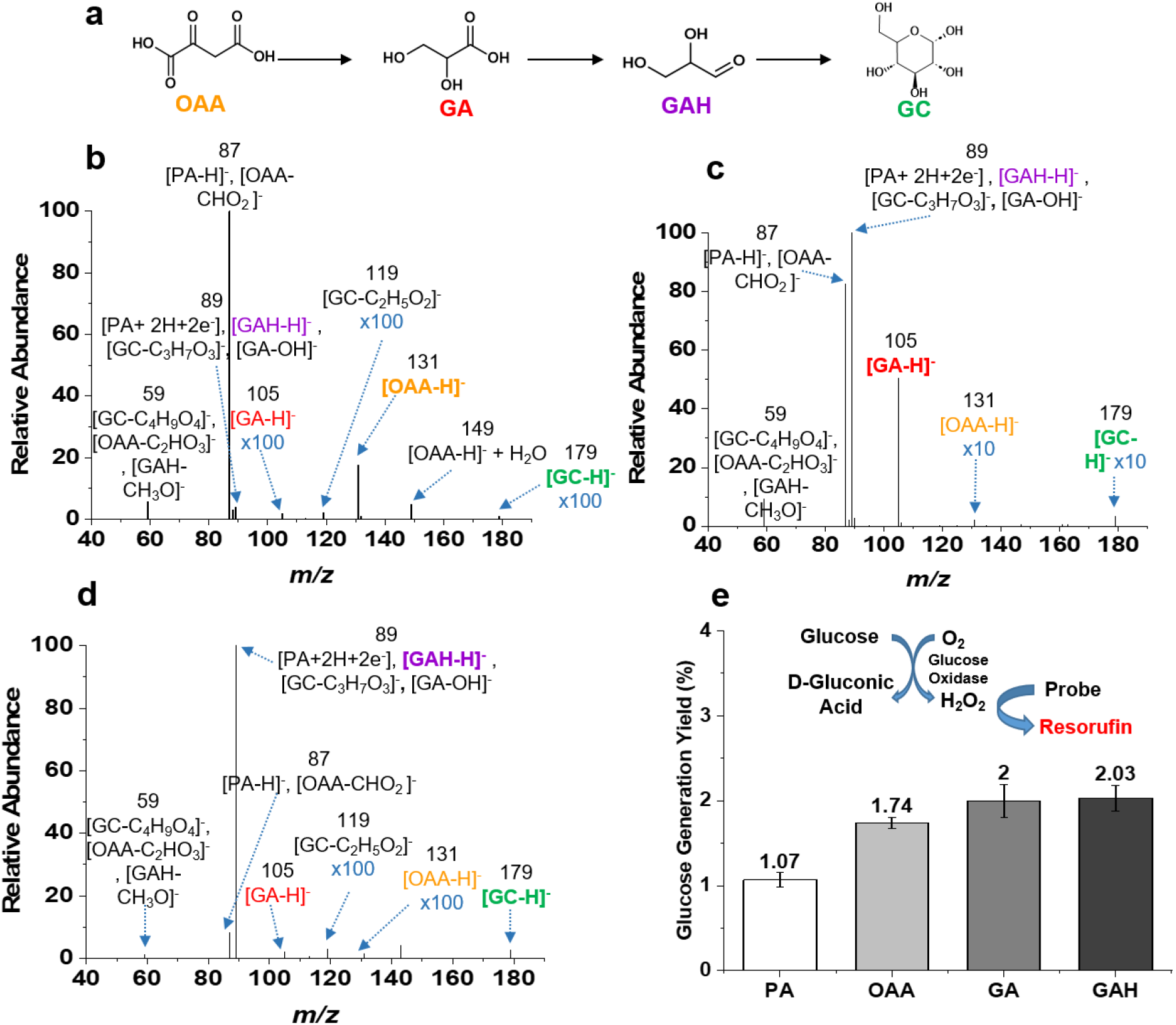
Mass spectrometric analysis of glucose generation from intermediates of gluconeogenesis. **a** Scheme of the reaction pathway. To verify the formation of glucose from gluconeogenic intermediates, including oxaloacetate (OAA), glycerate (GA), and glyceraldehyde (GAH), aqueous solutions containing each intermediate was sprayed and the products were mass spectrometrically recorded. All intermediates are color-coded for clarity. **b**-**d** Mass spectra recorded from the nebulized samples of each 10 μM solution of OAA, GA, and GAH into the MS inlet. **e** Glucose assay results showing glucose generation yield from each intermediate. A vibrating mesh generator was used to generate aqueous microdroplets. Error bars represent one standard deviation (SD) from three independent measurements.

### Bicarbonate-mediated conversion from pyruvate to oxaloacetate in microdroplets confirmed by ^13^C labeling

In biological gluconeogenesis, pyruvate is carboxylated to oxaloacetate using bicarbonate ion (HCO_3_-) as the carbon source. This reaction is catalyzed by pyruvate carboxylase and requires ATP and biotin as cofactors^50^. In our study, the formation of oxaloacetate was observed during glucose generation in aqueous microdroplets We hypothesized that glucose generation in microdroplets begins with the conversion of pyruvate to oxaloacetate through bicarbonate incorporation, analogous to biological gluconeogenesis but occurring without enzymes, ATP, or cofactors.

To examine the incorporation of bicarbonate into this reaction, we utilized an isotope-labeled carbon source, NaH^13^CO_3_ **(Fig. 4a)**. The control group consisted of degassed water nebulized with N_2_ gas. In this control experiment, no ^13^C-labeled oxaloacetate was detected (**Fig. 4b**). In contrast, in the experimental group with NaH^13^CO_3_, ^13^C-labeled oxaloacetate was observed (*m/z* 132.00124, –5.38 ppm) (**Fig. 4c**). Although the use of ^13^C-labeled bicarbonate enabled tracking of carbon incorporation, trace amounts of ^12^C-bicarbonate could not be fully excluded due to unavoidable atmospheric CO_2_ exposure under ambient conditions. These results demonstrate that HCO_3_− serves as the carbon donor in the formation of oxaloacetate within aqueous microdroplets, paralleling the carboxylation step of biological gluconeogenesis.

**Fig. 4.**
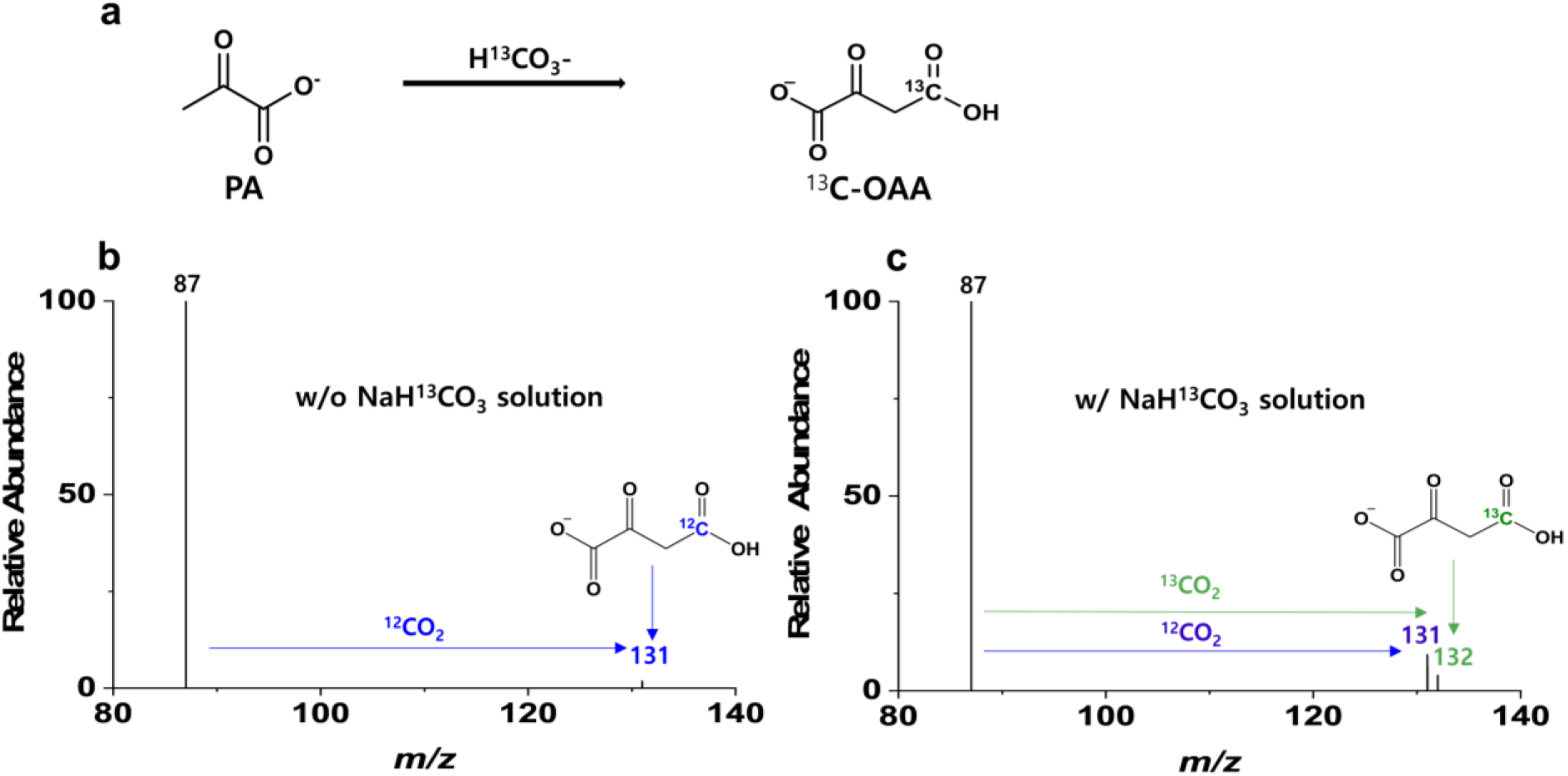
Identification of oxaloacetate intermediates from pyruvate aqueous microdroplets. **a** Experimental setup using NaH^13^CO_3_ as an isotopically labeled bicarbonate source A 100 μM solution of NaH^13^CO_3_ was prepared in H2O to examine the reductive carboxylation of pyruvate. **b** Mass spectrum of pyruvate-containing aqueous microdroplets generated without NaH^13^CO_3_. **c** Mass spectrum of pyruvate-containing aqueous microdroplets generated with NaH^13^CO_3_. The results confirm the bicarbonate-mediated reductive carboxylation of pyruvate to oxaloacetate in microdroplets.

### Characterization and kinetics of glucose generation in aqueous microdroplets

To investigate the optimal conditions for glucose generation in aqueous microdroplets, we measured the glucose generation yields of glucose with different pyruvate concentrations and droplet sizes. We first examined the dependence of glucose generation yield on reactant (pyruvate) concentrations, while keeping other parameters of inner capillary diameter, and N_2_ nebulization gas pressure constant at 50 μm and 0.8 MPa, respectively. Generation yield was quantified using a calibration curve based on standard glucose (R^2^ = 0.9987) **(Supplementary Fig. 6a)**. Higher glucose production yields were observed at lower pyruvate concentration **(Fig. 5a)**.

**Fig. 5.**
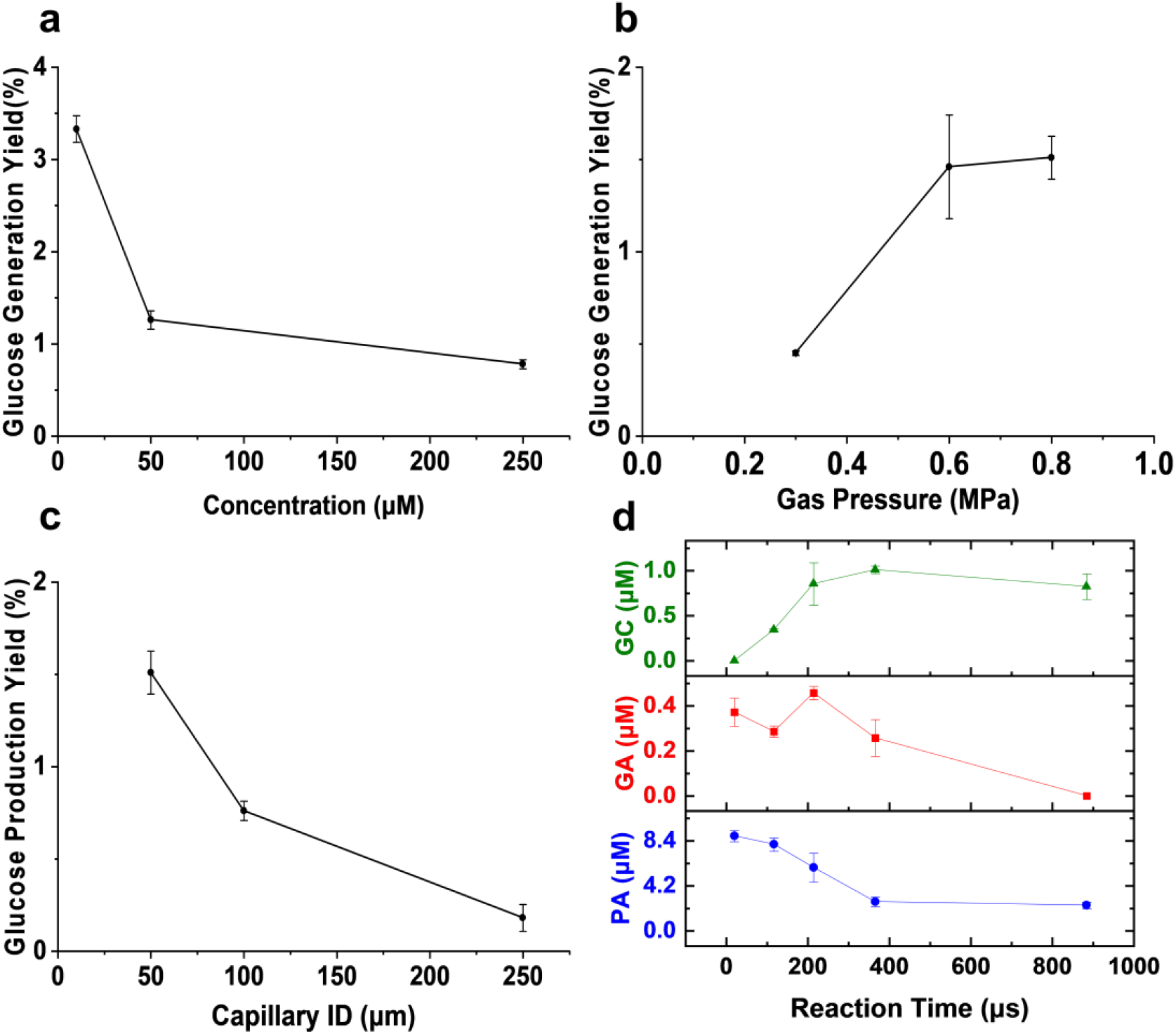
Characterizations and kinetics of glucose generation in pyruvate-containing aqueous microdroplets. **a** Dependence of glucose yield on pyruvate concentration. **b** Dependence on nebulizing N2 gas pressure. **c** Dependence on capillary inner diameter. **d** Time-dependent concentrations of pyruvate, glycerate, and glucose produced from microdroplets generated by spraying a 10 μM pyruvate solution. Reaction time was controlled by varying the distance between the capillary tip and the MS inlet. Error bars represent one standard deviation (SD) from three independent measurements.

**Fig. 6.**
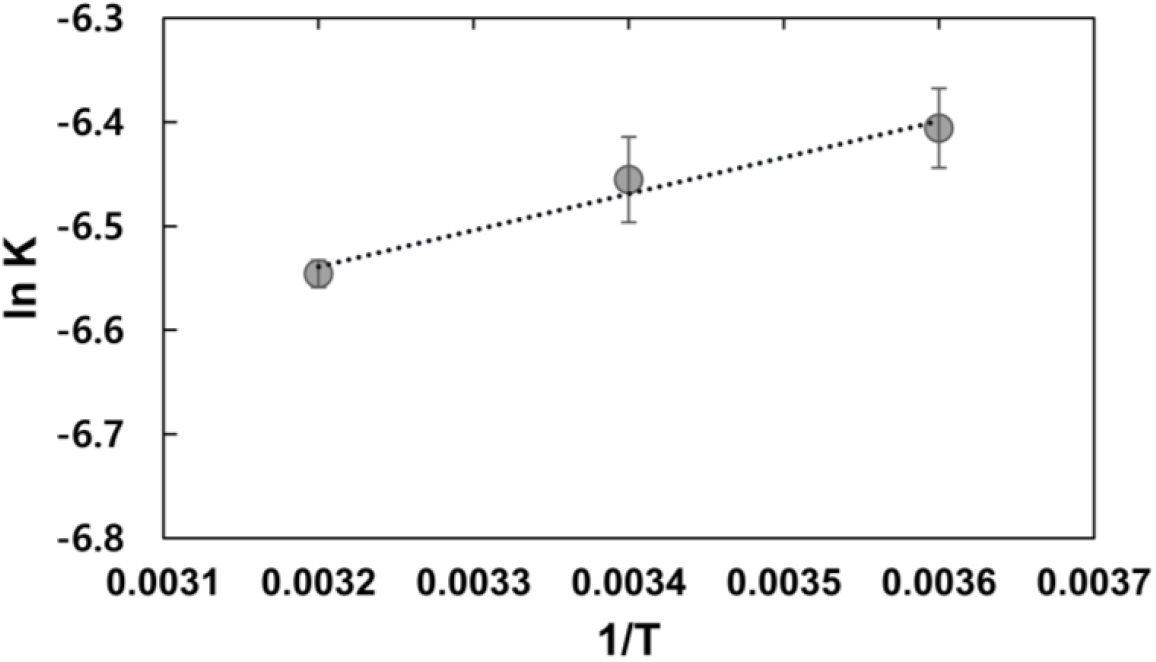
van’t Hoff plot for glucose generation from pyruvate in aqueous microdroplets. A 10 μM pyruvate solution was sprayed using an aerosol generator and glucose concentration were measured with a glucose assay. Thermodynamic parameters were calculated as Δ*H* = –0.70 kcal/mol and Δ*S* = – 0.015 kcal/mol·K. The slope of the plot corresponds to –Δ*H*/*R*, and the intercept to Δ*S*/*R*. Error bars represent one standard deviation (SD) from three independent measurements.

To evaluate the effect of droplet size on glucose production yield, we measured the production yield in different sizes of microdroplets generated by varying capillary inner diameters and nebulization gas pressure. Because the highest glucose generation yield was observed with 10 μM pyruvate solution **(Fig. 5a)**, we used a 10 μM pyruvate solution in all subsequent experiments. Smaller capillary inner diameter^41^ and higher gas pressure^51^ produced smaller microdroplets. Glucose yield increased with decreasing capillary diameter and increasing gas pressure **(Fig. 5b, c)**, indicating that microdroplets with higher surface-to-volume ratios enhance glucose production ^29^. This trend was also observed for reaction intermediates, including oxaloacetate **(Supplementary Fig. 7)**, glycerate **(Supplementary Fig. 8)**, and glyceraldehyde **(Supplementary Fig. 9)**. These results collectively suggest that the formation predominantly occurs at the microdroplet interface.

Then, we investigated the kinetics of glucose formation in aqueous microdroplet. A 10 μM pyruvate solution was nebulized, and the reaction time was regulated by adjusting the travel distance between the MS inlet and the capillary tip. Concentrations of pyruvate, glycerate, and glucose were quantified using calibration curves generated from standard reagents **(Supplementary Fig. 6a, b, d). Fig. 5d** presents the time-dependent concentration profiles of the reactant (pyruvate), central intermediate (glycerate), and reaction product (glucose). As reaction time increased, pyruvate concentration gradually decreased. Glycerate concentration initially increased, then declined to nearly zero at longer reaction times. Glucose concentration rose steadily and reached a maximum at 366 μs, followed by a slight decrease. Overall, these results confirm that glucose formation from pyruvate in aqueous microdroplets proceeds via a glycerate intermediate.

### Thermodynamics of glucose generation in aqueous microdroplets

Gluconeogenesis is an anabolic procedure in nature and does not proceed without a series of enzymes and ATPs. In contrast, we demonstrated that glucose formation in aqueous microdroplets occurs without any catalysts or energy input such as ATP. Therefore, the thermodynamic profile of glucose generation in aqueous microdroplets is expected to differ from that of biological gluconeogenesis. To investigate this, we calculated the Gibbs free energy change (ΔG) for glucose formation from pyruvate in aqueous microdroplets using the standard thermodynamic equation: Δ*G* = –*RT* ln *K*, where *R* is the universal gas constant and *K* is the equilibrium constant. Sprayed samples from aerosol generator were collected at a distance of approximately 100 mm from nozzle to the collection vial. At this distance, the reaction reached an equilibrium **(Fig. 5d)**. Δ*G* values were determined at three different temperatures. At room temperature, Δ*G* value was measured to be 3.78 kcal/mol **(Table 1)**. Additionally, we examined how temperature affects the thermodynamics of glucose formation. Δ*G* values increased slightly from 3.46 kcal/mol to 3.96 kcal/mol as temperature increased from 277 K to 310 K. From the van ‘t Hoff plot of the conversion reaction from pyruvate to glucose in aqueous microdroplets **(Fig. 6)**, the enthalpy and entropy value were determined to be Δ*H* = -0.70 kcal/mol, and Δ*S* = -0.015 kcal/mol?K. Although the calculated Δ*G* values of this reaction remain slightly positive, these results indicate that the microdroplet environment facilitates glucose formation by lowering the effective energetic barrier, enabling the reaction to proceed under ambient conditions.

**Table 1.** Calculated Gibbs free energy (ΔG) values for glucose generation in aqueous microdroplets at different temperatures.

| Temperature (K) | $\Delta G$ (kcal/mol) |
| --- | --- |
| 277 | 3.46 |
| 298 | 3.78 |
| 310 | 3.96 |

### Generation yields of different gluconeogenic intermediates in aqueous microdroplets

We then measured the conversion yields of various intermediates involved in glucose formation from aqueous microdroplets. Calibration curves for the quantitative analysis of pyruvate, oxaloacetate, glycerate, glyceraldehyde, and glucose were obtained using mass spectra of standard reagents across a concentration range of 0.001–100 μM (**Supplementary Fig. 6a–e**). A reactant concentration of 10 μM was chosen because glucose yield showed an inverse correlation with reactant concentration (**Fig. 5a**). The N2 gas pressure was maintained at 0.8 MPa, and the microdroplet travel time was set to approximately 210 μs.

The conversion yields of each intermediate are summarized in **Table 2**. The glucose yield from pyruvate-containing aqueous microdroplets was 3.33%. The highest glucose yield, 4.68%, was obtained from glyceraldehyde. Previous studies have reported the reduction of pyruvate to lactate ^33^. Notably, glyceraldehyde and lactate share the same molecular formula (C_3_H_6_O_3_). To accurately quantify glyceraldehyde formation from each precursor, including pyruvate, oxaloacetate, and glycerate, we used calibration curves based on the absolute intensities of their characteristic fragment ions, as described previously (**Supplementary Figs. 10 and 11**)^52^. In summary, these results confirm that both intermediates and glucose were generated in aqueous microdroplets from all four reactants, namely pyruvate, oxaloacetate, glycerate, and glyceraldehyde, with yields ranging from 2.14 to 7.2 %.

**Table 2.** The conversion yields of aqueous microdroplets reactions.

| Reactant concentration:<br>10 μM | Species detected in microdroplets (%) | | | | |
| --- | --- | --- | --- | --- | --- |
|  | 1 (PA) | 2 (OAA) | 3 (GA) | 4<sup>a</sup> (GAH) | 5 (GC) |
| 1 (PA) | - | 7.2±0.45 | 2.34±0.22 | 2.51±0.28 | 3.33±0.15 |
| 2 (OAA) | 26.25±3.96 | - | 2.14± 0.09 | 4.42±0.82 | 2.72±0.06 |
| 3 (GA) | 6.79±1.34 | 57.71±6.76 | - | 6.18±0.51 | 2.98±0.06 |
| 4 (GAH) | 0 | 13.88±3.10 | 22.32±2.93 | - | 4.68±0.15 |

<sup>a</sup> Glyceraldehyde and lactate share an identical $m/z$ of 89. To distinguish between them, the ratios of the absolute intensities of their main fragment ions were analyzed using compound-specific calibration curves (**Supplementary Fig. 10a, b**). Yields are reported as mean ± standard deviation ( $n = 3$ ).

## Discussion

Microdroplet chemistry has emerged as a remarkable platform for accelerating chemical transformations and promoting the synthesis of organic compounds^29,53^. Here, we demonstrate that glucose can be synthesized abiotically from pyruvate in aqueous microdroplets, without the need for enzymes, catalysts, or added energy sources. The reaction proceeds within a sub millisecond timescale under ambient conditions. We also detect key intermediates shared with biological gluconeogenesis, including oxaloacetate, glycerate, and glyceraldehyde.

Glucose production on Earth today is dominated by photosynthesis, but its origins before the evolution of photosynthetic organisms remain unresolved. Despite the centrality of glucose in metabolism, its formation under non-enzymatic conditions has not been clearly established. Our results provide experimental support for glucose synthesis from a simple three carbon precursor in aqueous microdroplets, suggesting a possible prebiotic route to this fundamental sugar.

Aqueous microdroplets, naturally present in aerosols, clouds, and sea spray, would have been abundant in the early stage of the Earth^54^. Even in the absence of enzymes and photosynthetic organisms in the early stage of the Earth, these aqueous microdroplets may play a role as a natural catalyst in prebiotic conditions to enable the assembly of energetically costly compound, glucose, from simple C_3_ compound, pyruvate. Previous studies have proposed that monosaccharides could arise from small carbon species such as formaldehyde, glycolaldehyde, or glyceraldehyde ^55,56^. By contrast, we show that pyruvate, both chemically simple and metabolically central, can serve as a direct precursor to glucose in a catalyst free system. This may help to understand how energy fuel of life can be made in prebiotic conditions.

Although previous studies in microdroplet chemistry have revealed accelerated reactivity and non-enzymatic synthesis of diverse biomolecules^29,32,33,53,57,58^, the direct formation of a six carbon sugar ring structure has remained elusive. Here we show that glucose—a cyclic six carbon compound essential to modern metabolism—can be assembled from a simple three carbon precursor, pyruvate, without the need for catalysts, enzymes, or external energy sources. This expands the known synthetic capabilities of microdroplet systems and establishes a new paradigm for sugar formation in abiotic environments.

Our findings uncover a chemically plausible, catalyst free route to glucose synthesis under ambient conditions. This demonstrates how microdroplets can act not only as physical compartments but also as functional chemical reactors capable of mediating complex carbon transformations. While the implications for prebiotic chemistry are evident, the broader significance lies in establishing aqueous microdroplets as dynamic and sustainable platforms for the green synthesis of structurally complex organic molecules, with potential relevance to environmental chemistry, industrial bioprocessing, and pharmaceutical innovation.

## Acknowledgements

This research was supported by Basic Science Research Program through the National Research Foundation of Korea (NRF) funded by the Ministry of Education(2022R1A6A1A03063039), the National Research Foundation of Korea(NRF) grant funded by the Korea government (MSIT)(2023R1A2C200723812), and Creative-Pioneering Researchers Program through Seoul National University.

