## Supplementary Information for "Gluconeogenesis in Aqueous Microdroplets: Non-Enzymatic Generation of Glucose"

**full $^1$H NMR spectral data of sprayed pyruvate and standard glucose**

**Standard pyruvate**: $^1$H NMR (400 MHz, $D_2O$) δ 2.41 (s), 1.68 (s), 1.57 (s).

**Sprayed pyruvate**: $^1$H NMR (400 MHz, $D_2O$) δ 5.63 (s), 4.79 (s), 4.19 (d, *J* = 6.4 Hz), 3.91 (d, *J* = 1.9 Hz), 3.71 (s), 3.59 (d, *J* = 12.0 Hz), 2.36 (s), 1.64 (s), 1.54 (s).

**Standard glucose**: $^1$H NMR (400 MHz, $D_2O$) δ 5.22 (s), 4.79 (s), 3.90 (d, *J* = 2.2 Hz), 3.87 (d, *J* = 2.2 Hz), 3.70 (s), 3.53 (d, *J* = 3.8 Hz).

### Supplementary method

#### Quantification of glyceraldehyde and lactate

Standard reagents were diluted in $H_2O$ (0.01-1 μM) and nebulized to a MS inlet. Higher-energy collisional dissociation (HCD) voltage at 15 V was used for molecular identification and quantitation. Pyruvate reduces into lactate and also converts into glyceraldehyde. Although lactate and glyceraldehyde are isomers that share the same nominal mass (*m/z* 89), their most abundant fragment ions differ: m/z 71 for glyceraldehyde and m/z 87 for lactate (**Supplementary Fig. 4a, Supplementary Fig. 10**). To deconvolute the overlapping signal at m/z 89, the relative contributions of each compound were calculated based on the intensities of their diagnostic fragment ions, using previously established methods[1].

The intensities of *m/z* 71 and 87 in the HCD spectra of microdroplet reaction products acquired by spraying 10 μM pyruvate is denoted as *M71* and *M87*, respectively. Because the HCD tandem MS/MS spectrum for glyceraldehyde and lactate exhibits their own unique ratios between *M71* and *M87*, the relative abundance between glyceraldehyde and lactate can be estimated by calculating the relative contributions to each fragment ion intensity in MS/MS spectra.

Because the unique ratio between *m/z* 71 and *m/z* 87 in the MS/MS spectrum of standard glyceraldehyde is 1 to 0.47, the relative abundance of the indicator ion of glyceraldehyde at *m/z* 71 and *m/z* 87 can be set as $x$ and 0.47*$x$, respectively. Similarly, the unique ratio between *m/z* 71 and *m/z* 87 in the MS/MS spectrum of standard lactate is 1 to 1.82, the relative abundance of the indicator ion of lactate at *m/z* 71 and *m/z* 87 can be set as $y$ and 1.82*$y$, respectively.

The following equation group presents the sum of the intensities for glyceraldehyde and lactate for each *M71* and *M87* (equation (1) and (2)).

$$x + y = M71 \quad (1)$$

$$0.47x + 1.82y = M87 \quad (2)$$

The values for *M71* and *M87* are obtained from the tandem MS/MS spectrum of sprayed microdroplet reaction products. Then the concentration of glyceraldehyde and lactate can be determined using the quantitation curves **(Supplementary Fig. 11)**.

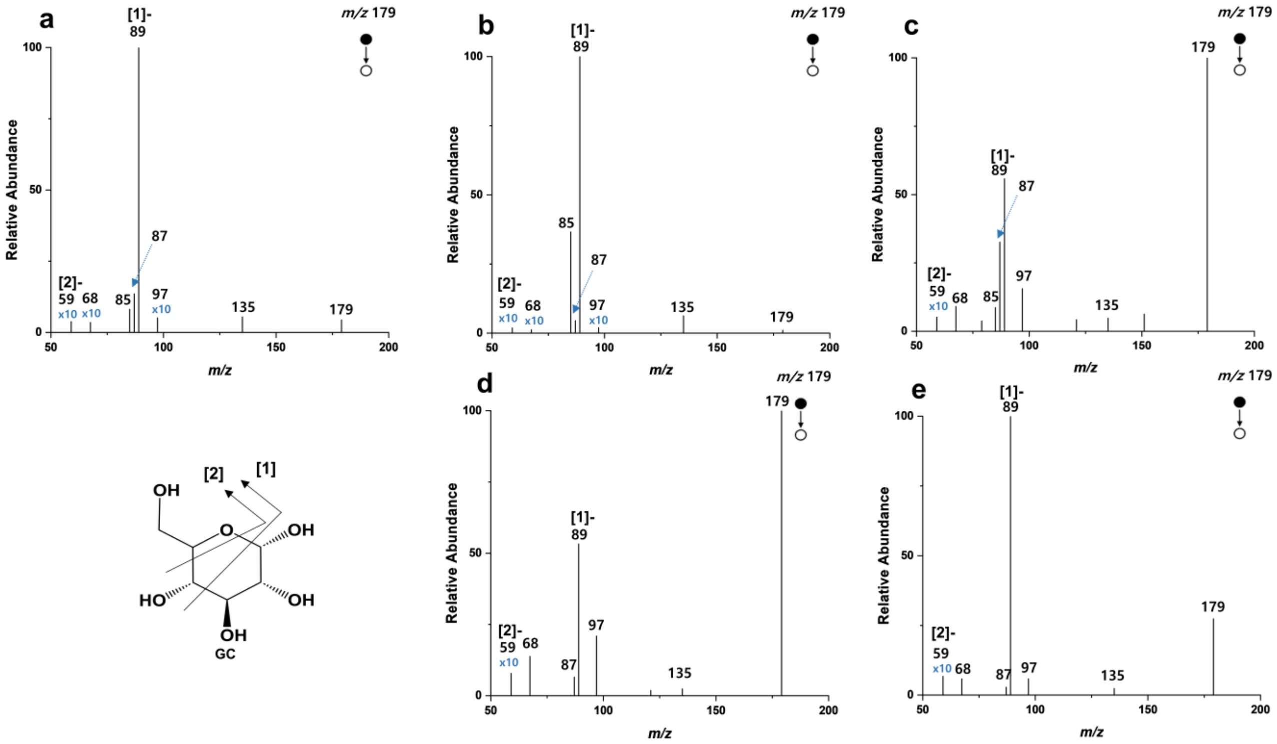

**Supplementary Fig. 1** Molecular identification of glucose generated by spraying aqueous microdroplets containing (**b**) pyruvate, (**c**) oxaloacetate, (**d**) glycerate, and (**e**) glyceraldehyde, using tandem MS/MS spectra. **a** MS/MS spectra of standard glucose**; (b–e)** MS/MS spectra of products generated from sprayed pyruvate, oxaloacetate, glycerate, and glyceraldehyde, respectively.

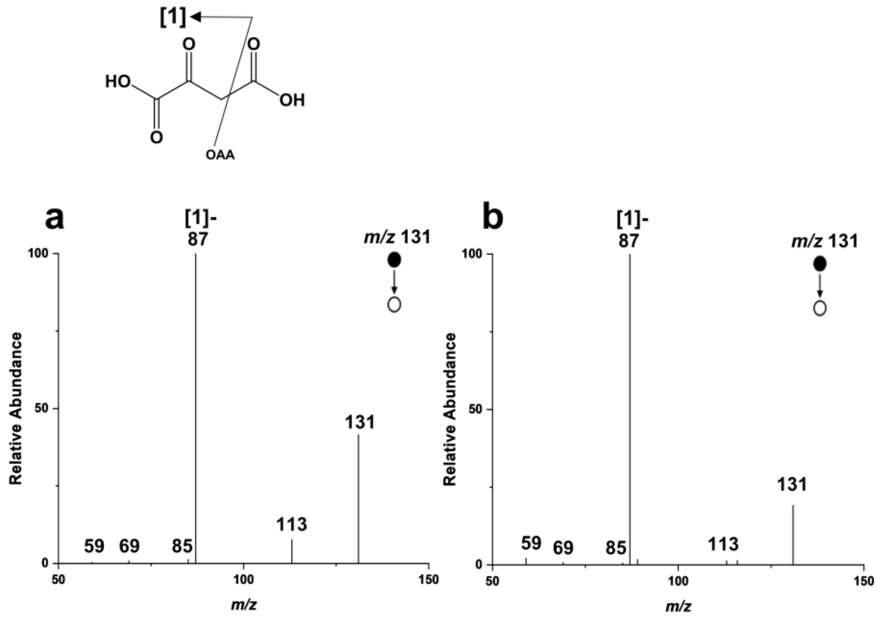

**Supplementary Fig. 2** Molecular identification of oxaloacetate generated by spraying aqueous microdroplets containing **b** pyruvate using MS/MS spectra. **a** Standard oxaloacetate. **b** Sprayed pyruvate.

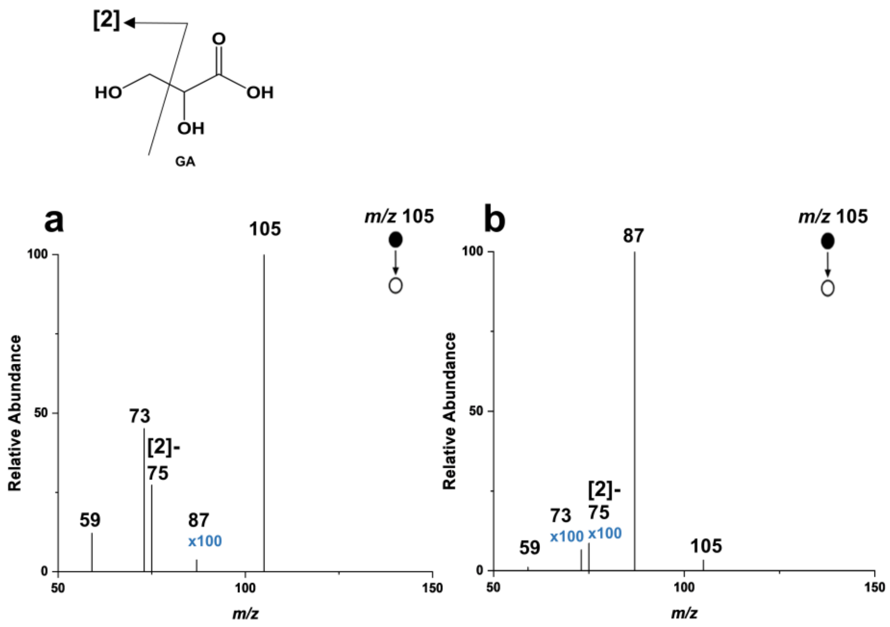

**Supplementary Fig. 3** Molecular identification of glycerate generated by spraying aqueous microdroplets containing **b** pyruvate using MS/MS spectra. **a** Standard glycerate. **b** Sprayed pyruvate.

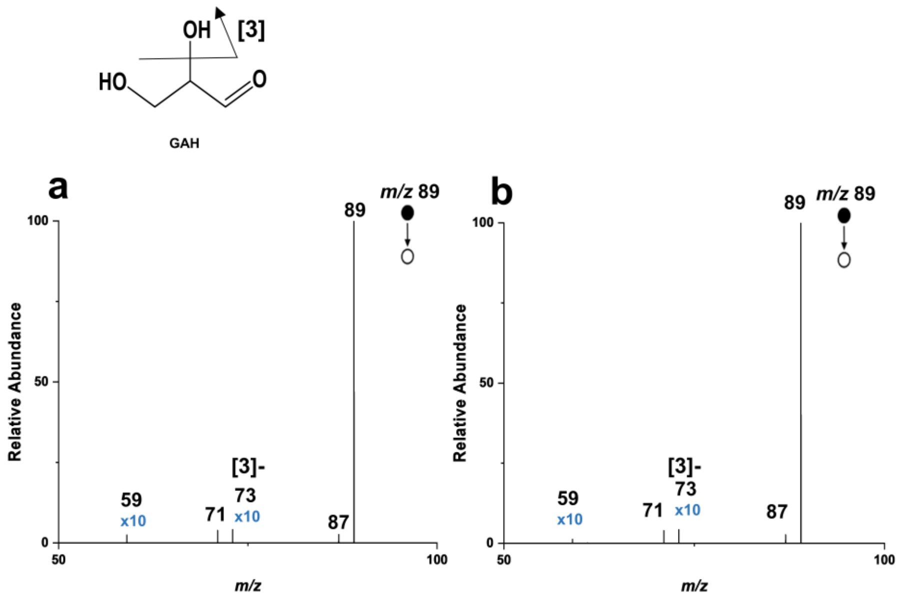

**Supplementary Fig. 4** Molecular identification of glyceraldehyde generated by spraying aqueous microdroplets containing **b** pyruvate using MS/MS spectra. **a** Standard glyceraldehyde. **b** Sprayed pyruvate.

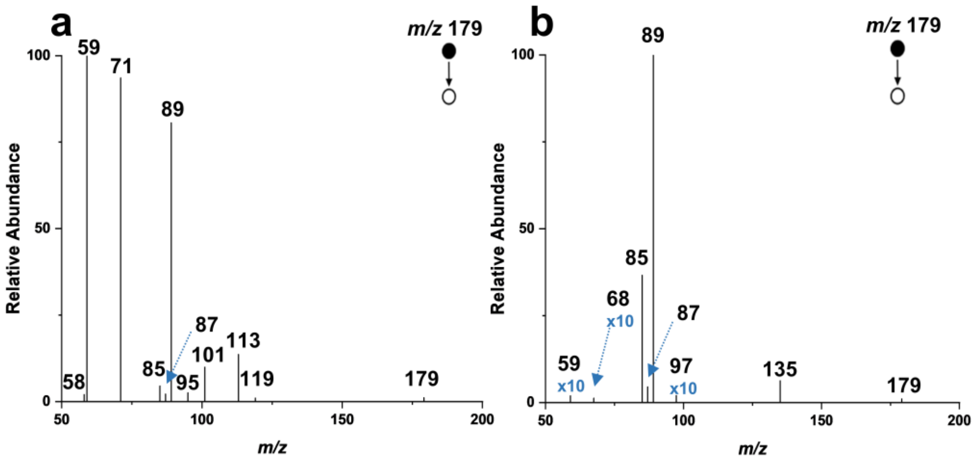

**Supplementary Fig. 5** Conventional gluconeogenesis pathway includes fructose. Fructose and glucose have the same *m/z* 179. Different MS/MS spectrum between **a** standard fructose and **b** products from microdroplets shows an exclusion of fructose.

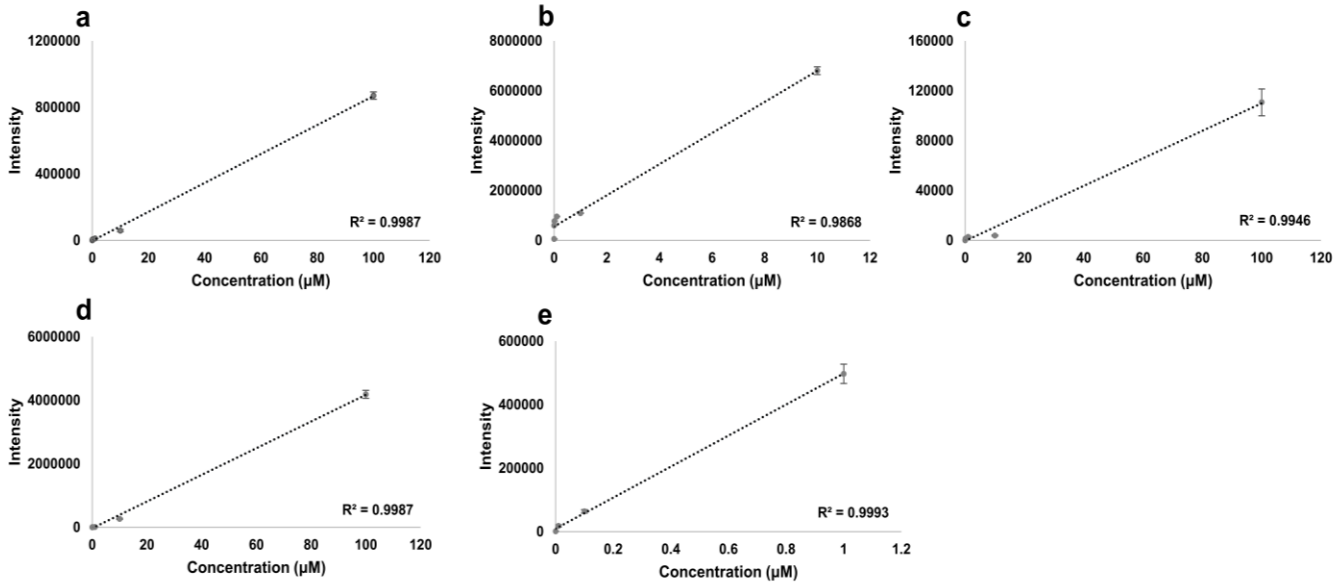

**Supplementary Fig. 6** Quantitation curves for **a** glucose, **b** pyruvate, **c** oxaloacetate, **d** glycerate, and **e** glyceraldehyde. Standard reagents were diluted in $H_2O$ (0.001-100 µM) and each quantitation curves were acquired using mass spectrometry. Error bars represent one SD from triplicate measurements.

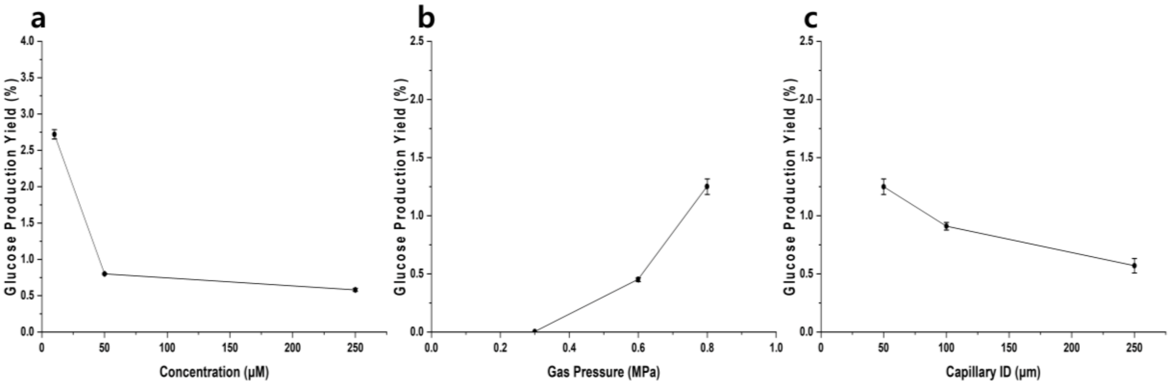

**Supplementary Fig. 7** Effect of **a** oxaloacetate concentration, **b** $N_2$ nebulizing gas pressure, and **c** capillary I.D. on glucose generation yield. Error bars represent one SD from triple measurements.

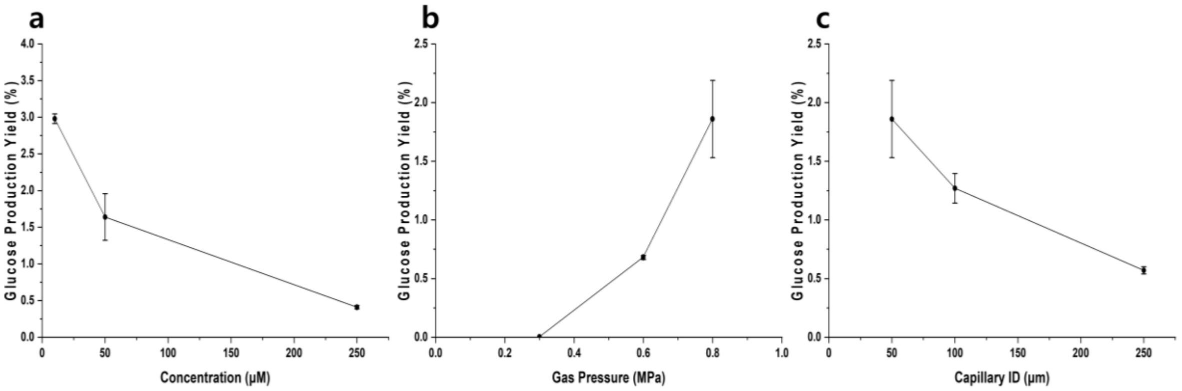

**Supplementary Fig. 8** Effect of **a** glycerate concentration, **b** $N_2$ nebulizing gas pressure, and **c** capillary I.D. on glucose generation yield. Error bars represent one SD from triple measurements.

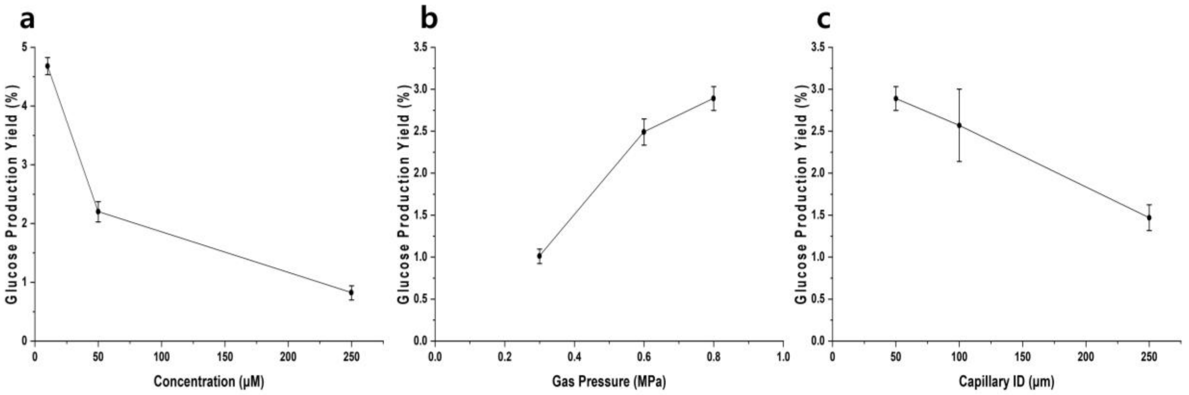

**Supplementary Fig. 9** Effect of **a** glyceraldehyde concentration, **b** $N_2$ nebulizing gas pressure, and **c** capillary I.D. on glucose generation yield. Error bars represent one SD from triple measurements.

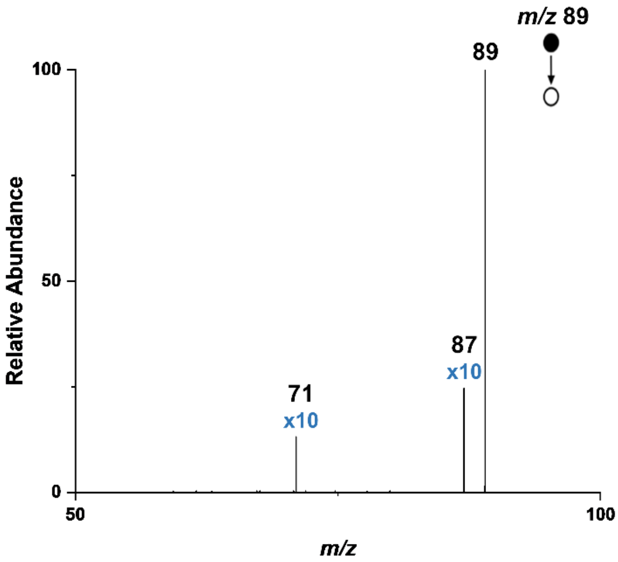

**Supplementary Fig. 10** MS/MS spectrum of standard lactate

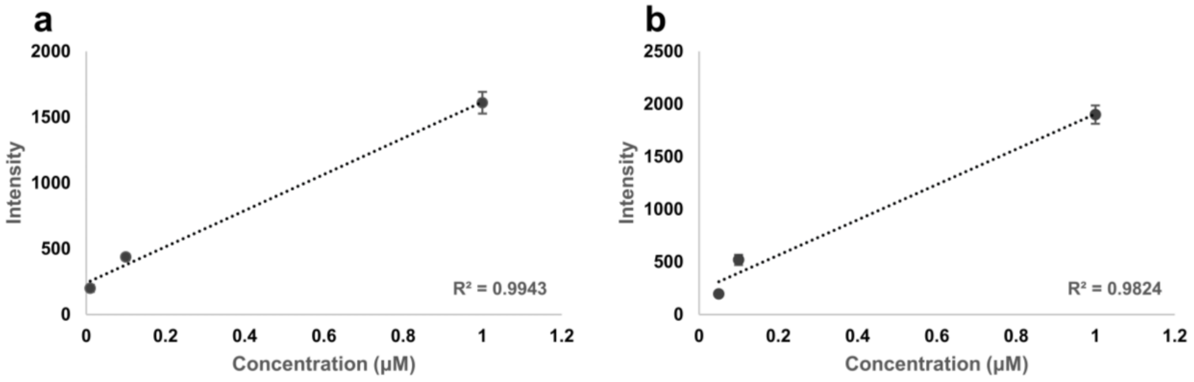

**Supplementary Fig. 11** Quantitation curves of **a** glyceraldehyde and **b** lactate.
